# A multi-tissue RNA-seq atlas of adult medaka (*Oryzias latipes*) under salinity and winter–summer sampling contrasts

**DOI:** 10.64898/2026.10.27.762344

**Authors:** Takafumi Katsumura, Yusuke Takehana, Atsushi Toyoda, Kiyoshi Naruse

**Affiliations:** Department of Human Life Design and Science, Faculty of Design, Kyushu University, Fukuoka, Japan; Graduate School of Bioscience, Nagahama Institute of Bio-Science and Technology, Nagahama, Japan; Department of Biological Sciences, Faculty of Advanced Science and Technology, Kumamoto University, Kumamoto, Japan; Advanced Genomics Center, National Institute of Genetics, Mishima, Japan; Comparative Genomics Laboratory, National Institute of Genetics, Mishima, Japan; Laboratory of Bioresources, National Institute for Basic Biology, Okazaki, Japan

**Author notes:** **Corresponding author:** Takafumi Katsumura.

## Abstract

Medaka (*Oryzias latipes*) is a euryhaline and eurythermal teleost widely used in genetics and environmental biology. We generated an RNA-sequencing atlas of 11 adult tissues (brain, eye, gill, heart, intestine, kidney, liver, muscle, spleen, ovary, testis) of the strain d-rR/TOKYO, sampled under two independent environmental contrasts: salinity (freshwater versus seawater) and season (winter versus summer). The freshwater and summer groups, reared indoors and outdoors under long days, provide a third contrast between laboratory and outdoor rearing. Typically, one male and one female were sampled per condition, with ten tissues taken from each individual. Ninety libraries were quantified against the medaka RefSeq transcript set (GCF_053564925.1). The resource comprises gene-level TPM and count matrices, differential expression tables for each tissue and contrast, tissue-specificity indices, human ortholog annotations, sample metadata, and a self-contained interactive browser. Because replication is limited and each condition was sampled on its own date, the differential expression tables are intended for hypothesis generation. The dataset supports studies of gene-expression plasticity and comparisons between medaka and humans.

## Background & Summary

The medaka (*Oryzias latipes*) is a well-established vertebrate model for genetics, development, and environmental physiology. A reference genome assembly, transgenic and genome-editing tools, and community resources such as the National BioResource Project support its use^1–3^. Wild populations are also accessible: across East Asia the species forms genetically distinct groups, and within the Japanese archipelago local populations range from Okinawa (26° N) to Aomori (40° N), inhabiting diverse environments and showing considerable phenotypic variation^4,5^. Medaka is unusual among model fishes^6,7^ in combining two forms of environmental tolerance: it is euryhaline (capable of surviving from freshwater to full-strength seawater^8^) and eurythermal (able to withstand temperatures from near 0 °C to over 30 °C^9^). This combination allows gene-expression plasticity, that is, how a single genome adjusts transcription across organs in response to environmental change, to be examined within a single species.

Salinity and season are two of the environmental factors that medaka contend with directly, each imposing demands across multiple organ systems^7^. Acclimation to changing salinity is a comprehensive physiological process: the gills remodel their ionocytes and ion-transport mechanisms, the kidneys modify water and ion regulation, and the intestines adjust their absorptive functions, such that a common osmotic challenge triggers distinct transcriptional responses in different organs^10–12^. Seasonal change, driven by covarying temperature and photoperiod, similarly involves numerous tissues: membrane lipids are remodeled to preserve functionality across temperature ranges, heat-shock and metabolic pathways are modulated, and reproductive timing aligns with favorable conditions^9,13^. Given that these responses are distributed throughout the organism, accurately capturing them necessitates a comprehensive assessment of the entire body.

These environmentally induced responses are also of evolutionary interest because phenotypic plasticity is increasingly seen as a starting point for adaptation. A trait induced in a new or recurring environment can be refined and eventually become genetically fixed, a process termed plasticity-led evolution^14–16^. Regulatory and epigenetic changes typically mediate the initial, reversible response, and subsequent genetic fixation proceeds through new mutations or selection on standing variation. We recently provided molecular evidence for this process in medaka, showing that the loss of DNA methylation sites abolishes an environmentally plastic trait, after which standing genetic variation is recruited and fixed through selection^17^. Seasonal plasticity in eye phototransduction and opsin expression modifies color perception and mating behavior in medaka^18^, and a transcriptional program underlies the circannual rhythm of gonadal development^19^. From this perspective, the plastic responses of gene expression elicited by the environment constitute the substrate on which selection acts, and cataloging these responses systematically provides a reference for studying their subsequent fixation.

Nevertheless, the environmental responses of medaka have been characterized in a fragmented manner, typically focusing on a single tissue, a single environmental factor, or one gene family at a time^9–12^. Multi-tissue transcriptome data do exist, but were generated to assemble gene repertoires and to construct gene models^20,21^, and in both cases the fish were maintained under constant laboratory conditions. These data are therefore not suited to examining how gene expression responds to the environment. Laboratory rearing also holds temperature and photoperiod constant, whereas outdoor conditions fluctuate daily, and in fish, acclimation to constant and to daily fluctuating temperatures leads to different gene-expression profiles^22^. How closely expression under standard laboratory rearing matches that under outdoor conditions therefore bears on how laboratory findings are extended to other settings^23^. In mammals, adult expression atlases are widely used in functional genomics^24,25^; among fishes, a comparable resource has recently been reported for killifish^26^. For medaka, however, a quantitative expression atlas of multiple tissues collected from individual fish under contrasting environmental conditions has not been available.

We generated an RNA-seq dataset covering 11 tissues of adult medaka sampled under two independent environmental contrasts: salinity (freshwater-versus-seawater) and season (a winter-versus-summer sampling contrast). These contrasts were sampled in separate groups of individuals, rather than combining both salinity and season treatments within the same fish (i.e., not a crossed salinity-by-season design). Because the freshwater group was reared indoors under constant conditions and the summer group outdoors, both under long days, the two groups also provide a third, cross-cohort contrast between laboratory and outdoor rearing. For each somatic tissue, we sampled typically one male and one female per condition; gonads were represented by a single sex. Collecting multiple tissues from each individual facilitates within-individual comparisons of expression patterns across tissues, although different individuals were sampled under each environmental condition and each condition was sampled on its own date.

After quality control, we quantified 90 libraries against the medaka RefSeq transcript set of assembly GCF_053564925.1^27^, providing gene-level expression annotated with gene symbol, gene name, and Entrez identifier. We release normalized expression matrices, sample principal component scores, per-axis, per-tissue differential expression tables, tissue-specificity indices, human ortholog annotations, sample metadata, and an interactive, self-contained browser, the Medaka RNA-seq Dashboard. The browser allows gene expression to be inspected by tissue and condition without programming. Genes are linked to their human orthologs to support translational and evolutionary work, building on medaka’s utility for analyzing genome– environment interactions^28^. Because biological replication per condition is limited, the differential-expression tables are provided as exploratory and hypothesis-generating rather than as definitive statistical tests. This dataset may support studies of expression plasticity, tissue-specific responses to osmotic, seasonal and rearing conditions, candidate gene prioritization, and, through human orthologs, comparative analyses relevant to vertebrate adaptation.

## Methods

### Ethics statement

All animal experiments were performed in accordance with the relevant institutional guidelines and were approved by the Institutional Animal Care and Use Committee of the National Institute for Basic Biology (approval no. 25A086) and by the Nagahama Institute of Bio-Science and Technology (approval no. 098).

### Animals and rearing

All fish were the medaka (*Oryzias latipes*) experimental strain d-rR/TOKYO (NBRP Strain ID: MT837). The two environmental contrasts were obtained from two separate fish cohorts (salinity and season).

For the salinity cohort, the fish were reared at the National Institute for Basic Biology (NIBB; Okazaki, Japan) under constant indoor conditions (25 °C, 14 hours of light and 10 hours of darkness) in either freshwater or full-strength artificial seawater (SEALIFE, Marinetech). The seawater fish were the offspring of parents that had been acclimated to seawater. They were fertilized, hatched and reared to sexual maturity in seawater, so they had never been exposed to freshwater. The specific gravity of the seawater was kept at approximately 1.02 and checked once per week with a hydrometer (Tetra). Each group of about 10 individuals was kept in one recirculating 9 L tank (Aquatic Habitats). The survival to sexual maturity in our seawater condition was almost 80%, similar to that in freshwater.

For the seasonal cohort, fish were maintained in a single outdoor rooftop tank (Sekisui Container S-54-II, 552 × 400 × 309 mm; approximately 20 fish) under natural conditions at the Nagahama Institute of Bio-Science and Technology (Nagahama, Japan). Sexually mature adults were sampled from the same tank in winter (22 December 2023, ∼13:00) and in summer (3 June 2024, ∼13:00). Water temperature was not recorded. As a reference, air temperature at the Nagahama station of the Japan Meteorological Agency (https://www.data.jma.go.jp/stats/etrn/index.php) was summarized over the 28 days up to and including each sampling date. The mean daily temperature was 7.0 °C in winter and 17.3 °C in summer, and the mean daily range (daily maximum minus daily minimum) was 8.5 °C and 10.7 °C, respectively. The day length was about 9.8 h on the winter sampling date and about 14.4 h on the summer sampling date (https://eco.mtk.nao.ac.jp/koyomi/index.html.en).

The two cohorts differ in how condition relates to tank. In the salinity cohort, each condition was kept in its own tank; the two tanks were of the same type and were held under the same temperature, photoperiod, and feeding regime. In the seasonal cohort, both samplings came from the same tank, so the fish were about five months older at the summer sampling, and fewer fish remained after the winter sampling.

### Sampling design and dissection

Fish were sexually mature adults; exact age and body size were not recorded. The fish were not fasted before sampling and were euthanized under eugenol anesthesia in accordance with the approved institutional protocols. Tissues were dissected in no fixed order, with care taken to avoid carry-over between tissues. The same procedure and the same tissue definitions were used for both cohorts. From each individual, ten tissue types were dissected and stored in RNA*later* (Thermo Fisher Scientific): whole brain (including the pituitary), whole eye, gill (entire branchial arches), heart, the entire digestive tract (esophagus to anus; operationally labeled ‘intestine’ herein), whole kidney (including both head and trunk kidney portions), liver (entire lobes), muscle (caudal trunk musculature without skin or bone), spleen (entire organ), and the gonad (ovary and testis). Because the gonad is sex-specific, the dataset covers eleven tissue types, of which each individual contributes ten (nine somatic tissues plus one gonad).

For each condition, one male and one female were sampled for the somatic tissues, and the gonads were represented by the corresponding single sex. Individuals are identified by condition (Fw, freshwater; Sw, seawater; Su, summer; W, winter), sex and a number; for example, SwM5 is seawater male #5. Library names add the prefix R (for RNA) and a tissue letter before the number; for example, RSwMT5 is the testis library of SwM5. Two additions to this scheme were retained. An additional seawater male (SwM5) was dissected on the same date as the other seawater fish (SwF2 and SwM1); its tissues were stored in RNA*later* at −80 °C, and RNA was extracted about eight months later, after the testis library of SwM1 had been excluded (see below). An additional winter female muscle RNA library (RWFM1) was prepared from a second winter female (WF1) to provide RNA-seq data matched to DNA from the same individual for another study. The sample composition by tissue and condition is summarized in Table 1; per-sample identifiers and attributes are provided in Supplementary Table S1. In total, 91 libraries were prepared; one library was excluded after quality control (see below), leaving 90 libraries for analysis.

**Table 1.** Sample composition by tissue and condition. Cells give the number of analyzed libraries by sex (M, male; F, female). For each somatic tissue, one male and one female were sampled per condition; gonads (ovary, testis) are single-sex, one individual per condition. Two additions increase specific cells: an additional seawater male (SwM5) raises all seawater somatic tissues to 2M+1F, and the muscle library of a second winter female (RWFM1, from WF1) raises winter muscle to 1M+2F. The freshwater and summer groups also form the rearing contrast. Counts are the 90 analyzed libraries; one seawater testis library (RSwMT1) was excluded after quality control as a mislabeled tissue (91 libraries were deposited). Per-sample identifiers, attributes, and processing batches are given in Supplementary Table S1.

| **Tissue** | **Salinity** |  | **Season** |  |
|---|---|---|---|---|
|  | **Freshwater** | **Seawater** | **Winter** | **Summer** |
| Brain | 1M+1F | 2M+1F | 1M+1F | 1M+1F |
| Eye | 1M+1F | 2M+1F | 1M+1F | 1M+1F |
| Gill | 1M+1F | 2M+1F | 1M+1F | 1M+1F |
| Heart | 1M+1F | 2M+1F | 1M+1F | 1M+1F |
| Intestine | 1M+1F | 2M+1F | 1M+1F | 1M+1F |
| Kidney | 1M+1F | 2M+1F | 1M+1F | 1M+1F |
| Liver | 1M+1F | 2M+1F | 1M+1F | 1M+1F |
| Muscle | 1M+1F | 2M+1F | 1M+2F | 1M+1F |
| Spleen | 1M+1F | 2M+1F | 1M+1F | 1M+1F |
| Ovary (♀) | 1F | 1F | 1F | 1F |
| Testis (♂) | 1M | 1M | 1M | 1M |
| **Total libraries** | **20** | **29** | **21** | **20** |

### Nucleic acid extraction

Tissues were kept in RNA*later* at −80 °C for several days to several weeks before extraction, except those of SwM5, from which RNA was extracted on 7 November 2024, about eight months after sampling. Total RNA and genomic DNA were co-purified from each tissue with the AllPrep DNA/RNA Mini Kit (Qiagen). Tissue was homogenized with a single 2-mm zirconia bead in 350 µL Buffer RLT containing 3.5 µL 2-mercaptoethanol using a ShakeMan 6 homogenizer (Bio Medical Science) (3,000 rpm, 30 s, repeated up to six times). Lysates were passed through an AllPrep DNA spin column (genomic DNA retained for a separate study), and RNA was purified from the flow-through on an RNeasy column with on-column DNase I digestion, then eluted in 50 µL RNase-free water. RNA concentration was measured using NanoDrop and Qubit fluorometry (Thermo Fisher Scientific), and RNA integrity was assessed using the TapeStation 4150 (Agilent). RNA was stored at −80 °C until library preparation.

### Library preparation and sequencing

RNA-seq libraries were prepared with the Illumina Stranded mRNA Prep, Ligation kit (poly(A)-selected, strand-specific) and sequenced on an Illumina NovaSeq 6000 instrument in paired-end mode (2×151 cycles). Sampling dates and sequencing batches for each condition are given in Table 2.

**Table 2.** Sample processing by condition. Sampling date, and sequencing batch (defined by shipment date to the sequencing facility; libraries shipped on the same date were sequenced together) for each condition. RNA was extracted several days to several weeks after sampling, except for seawater SwM5 (see Notes). Freshwater and most seawater libraries share one sequencing batch (2024-05-21). All summer libraries are in one batch (2024-06-18) whereas winter libraries are in earlier batches, apart from RWFM1. The freshwater and summer groups also form the rearing contrast. Counts are of collected libraries (n = 91); one seawater library (RSwMT1) was excluded after QC, leaving 90 analyzed libraries.

| **Condition** | **Axis** | **Sampling date** | **Sequencing batch (shipment date)** | **Number of libraries** | **Notes** |
|---|---|---|---|---|---|
| Winter | Season | 2023-12-22 | 2024-03-12 | 9 |  |
| Winter | Season | 2023-12-22 | 2024-05-21 | 11 |  |
| Winter | Season | 2023-12-22 | 2024-06-18 | 1 | RWFM1 (muscle of WF1) |
| Summer | Season, Rearing | 2024-06-03 | 2024-06-18 | 20 |  |
| Freshwater | Salinity, Rearing | 2024-04-16 | 2024-05-21 | 20 |  |
| Seawater | Salinity | 2024-03-13 | 2024-05-21 | 20 | SwF2 and SwM1 |
| Seawater | Salinity | 2024-03-13 | 2024-11-12 | 10 | SwM5; RNA extracted on 2024-11-07 |

### Read processing and transcript quantification

Raw paired-end reads were processed with fastp v0.23.2^29^ (fastp -3 -q 15 -n 5 -t 1 -T 1 -l 20). Adapters were removed by the default paired-end overlap detection. The 3′ end was quality-trimmed with a sliding window (-3), bases with a Phred score below 15 were counted as unqualified and reads exceeding the default unqualified fraction were discarded (-q 15), reads with more than five ambiguous (N) bases were removed (-n 5), and reads shorter than 20 bp were discarded (-l 20). One base was trimmed from the 3′ end of each mate (-t 1 -T 1) to remove the terminal sequencing cycle, which on Illumina 2×151-cycle chemistry can show elevated error and base-composition skew. A decoy-aware salmon index was built from the RefSeq transcript set of the medaka telomere-to-telomere assembly GCF_053564925.1^27^, using the entire genome as decoy and a k-mer length of 31. Trimmed reads were quantified with salmon v1.10.3^30^ against this index with automatic library-type detection (-l A), selective alignment (--validateMappings), and GC- and sequence-bias correction (--gcBias --seqBias). Transcript-level estimates were summarized to the gene level with tximport v1.34.0 (R v4.4.2)^31^ with countsFromAbundance = “no”. Gene-level counts are tximport-derived estimated counts from Salmon transcript quantification rather than alignment-derived integer read counts. For DESeq2 modeling, these estimated counts were imported with DESeqDataSetFromTximport, which rounds them to integers and uses the average transcript lengths from tximport as normalization offsets. Transcript-to-gene assignments and gene annotation (gene symbol, description, and Entrez Gene ID) were parsed from the RefSeq annotation (genomic.gff) of GCF_053564925.1, including coding and non-coding biotypes (mRNA, lnc_RNA, ncRNA, rRNA, tRNA, snoRNA, snRNA, miRNA, antisense_RNA, and related transcript types). Libraries flagged use = FALSE were excluded, yielding gene-level TPM and estimated-count matrices of 24,488 genes across 90 libraries.

### Quality control and outlier exclusion

Sample structure was first examined across all 91 libraries, using principal component analysis of log2(TPM + 1) for the 5,000 most variable genes, after retaining genes whose mean TPM across all libraries was at least 1, with centered, unscaled prcomp. To diagnose a sample lying outside its expected tissue cluster, each sample was correlated (Pearson) with per-tissue centroids and with all other individual samples, using the same gene set as the principal component analysis. Per-tissue centroids were computed from the remaining samples, excluding the sample under evaluation. After exclusion, the gene filter and the selection of variable genes were reapplied to the remaining 90 libraries, and Fig. 1 is based on this rebuilt matrix. One library labeled as testis (RSwMT1) correlated most strongly with intestine rather than gonad, and its nearest-neighbor sample was an intestine library (RSwMI1, r ≈ 0.94). Because its expression profile was inconsistent with the assigned tissue, it was flagged use = FALSE, and excluded from the rebuilt matrices and all downstream analyses. The complete raw read set, including this library, is retained in the data repository.

**Fig. 1.**
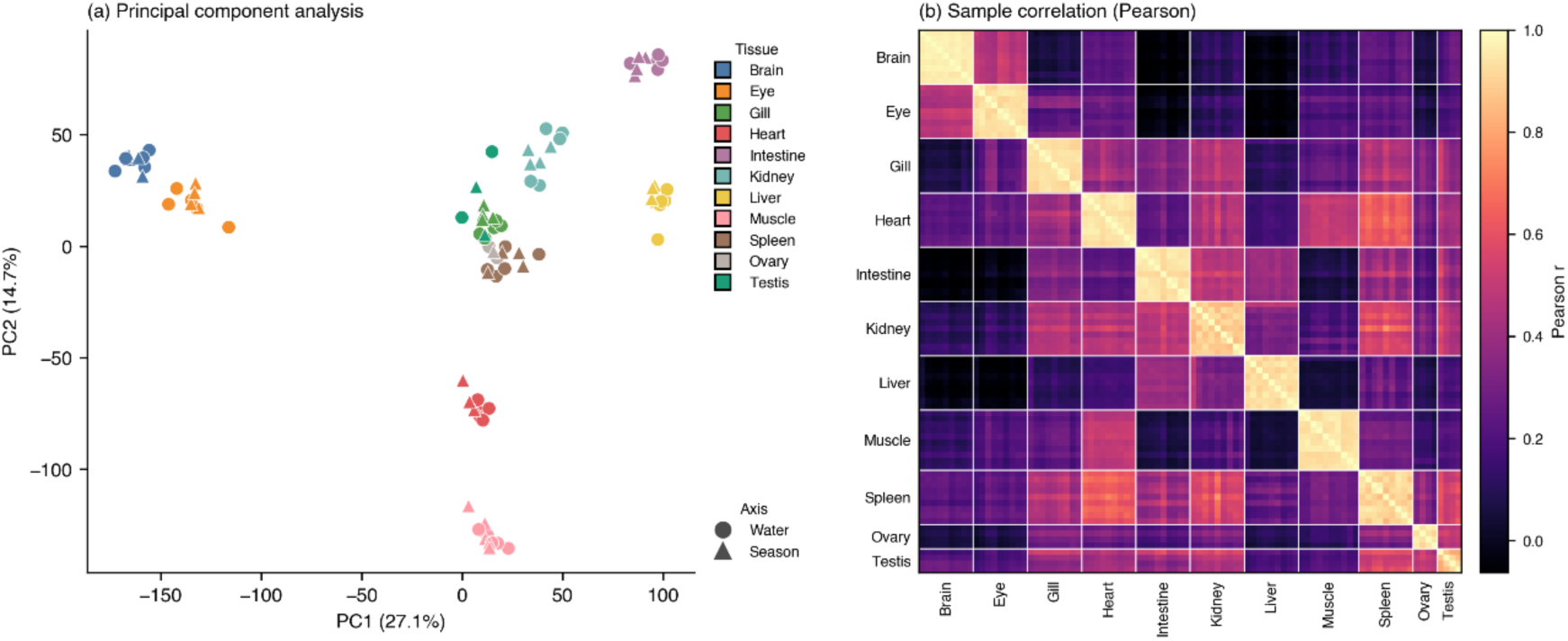
Sample structure. Both panels are computed from the same matrix: log2(TPM + 1) values for the 5,000 most variable genes among those with mean TPM ≥ 1 across the 90 analyzed libraries. (a) Principal component analysis (gene-centered, unscaled); points are libraries, colored by tissue and shaped by environmental axis. PC1 and PC2 account for 27.1% and 14.7% of the variance, respectively. (b) Pairwise Pearson correlation between libraries, computed across the same 5,000 genes. Libraries are ordered by tissue (in the order labeled) and, within a tissue, by sample identifier; no clustering method was applied. White lines delimit tissues.

### Differential expression analysis

Differential expression was assessed with DESeq2 v1.46.0^32^, separately for each tissue within each of three axes. The salinity axis contrasted seawater versus freshwater (reference: freshwater; positive log2 fold change indicates higher expression in seawater), and the seasonal axis contrasted winter versus summer (reference: summer; positive log2 fold change indicates higher expression in winter). A third axis (rearing) contrasted the summer group with the freshwater group (reference: freshwater group, labeled Lab; positive log2 fold change indicates higher expression in the summer group, labeled Out). Both groups were under long-day conditions (14 h light indoors; natural day length of about 14.4 h outdoors). They differ in facility, in sampling date and sequencing batch (Table 2), and in temperature (25 °C indoors; outdoors, a mean air temperature of 17.3 °C with a mean daily range of 10.7 °C over the 28 days before sampling). This axis reuses the freshwater and summer libraries and is therefore not independent of the other two. Only samples informative for the given axis were included.

For each tissue–axis combination, gene-level counts were modeled with the design ∼ Sex + condition when both sexes were present. Sex was included as a covariate to adjust for baseline sex-specific expression differences across the individual-level samples, yielding sex-adjusted condition effects. Sex-by-condition interactions were not modeled, because with one individual per sex and condition a model with an interaction term has no residual degrees of freedom in most comparisons. Gonads had a single individual per condition and were not tested. Genes were pre-filtered to those with at least 10 estimated counts in at least min(n₁, n₂) samples, where n₁ and n₂ are the per-condition sample sizes; combinations with fewer than two replicates per condition, or fewer than 100 genes passing the filter, were not tested. In total, 27 tissue-by-axis combinations were tested (nine somatic tissues on each of three axes). Log2 fold changes were shrunk with a normal prior (lfcShrink, type = “normal”). Genes were considered candidate genes meeting the exploratory DE threshold if they satisfied two criteria: control of the false discovery rate at 10% by the Benjamini–Hochberg procedure (adjusted p < 0.1), and a shrunken |log2 fold change| greater than 1.

The released tables report both the shrunken estimate and the unshrunken maximum-likelihood estimate, so users can apply either. Because biological replication per condition is limited and lacks independent tank-level replication, these differential-expression results are provided as exploratory, hypothesis-generating resources rather than powered statistical tests.

### Tissue-specificity index

A tissue-specificity index (τ)^33^ was computed for each gene across the eleven tissues. Per-tissue mean expression was obtained by averaging TPM over all retained samples of each tissue, and τ was calculated on log2(mean TPM + 1) as τ = Σ(1 − x^ᵢ)/(n − 1), where x^ᵢ is the per-tissue value divided by its maximum across the n = 11 tissues; τ ranges from 0 (ubiquitous) to 1 (tissue-restricted). The log2 transformation was applied to prevent extremely high-expressing genes from disproportionately dominating the specificity ranking. This log-scaled τ is a dataset-specific descriptive index and should not be directly interchanged with τ values calculated on untransformed expression values. Genes whose maximum per-tissue mean was below 1 TPM were considered undetected and assigned an undefined τ, because τ is invariant to overall expression level and would otherwise be dominated by genes near the detection limit. Each gene was additionally annotated with its most highly expressed (“enriched”) tissue. To check that the index is not driven by pooling the two cohorts, τ was also computed separately within the salinity cohort and within the seasonal cohort, using the same definition and the same 11 tissues. For each tissue, the top 10 genes with the highest τ were listed as a tissue-restricted gene set, restricted to genes with a maximum per-tissue mean expression of at least 100 TPM. The threshold was applied so that the listed genes are robustly detectable and therefore usable as markers.

### Human ortholog mapping

Medaka genes were mapped to human orthologs using the NCBI gene_orthologs dataset (https://ftp.ncbi.nlm.nih.gov/gene/DATA/gene_orthologs.gz, retrieved 2026-06-12; 18,040,422 pairwise ortholog records across all taxa), retaining pairs between *Oryzias latipes* (taxonomy ID 8090) and *Homo sapiens* (taxonomy ID 9606) in either orientation and de-duplicating on the medaka–human GeneID pair; human gene symbols were joined from the NCBI gene_info file for Homo sapiens (https://ftp.ncbi.nlm.nih.gov/gene/DATA/GENE_INFO/Mammalia/Homo_sapiens.gene_info.gz) on the human GeneID. For coverage statistics, “expressed” genes were defined as those with a maximum per-tissue mean ≥ 1 TPM (the same threshold used for the tissue-specificity index), and “tissue-enriched” genes as those with τ ≥ 0.8.

### Cross-species comparison of tissue specificity

To assess whether the released human-ortholog annotation supports cross-species comparison, tissue specificity was compared between medaka and a human reference. Human expression values were taken from the Human Protein Atlas (HPA) v25.1^34,35^ (rna_tissue_consensus.tsv, giving consensus normalized transcripts per million (nTPM) for each gene and tissue; retrieved 14 July 2026). The comparison was restricted to a shared nine-tissue panel: brain, heart, intestine, kidney, liver, muscle, spleen, ovary and testis. Eye and gill were excluded because the human panel has no corresponding tissue. HPA tissues were mapped onto this panel one-to-one, except that the individual brain regions listed in HPA v25.1 were averaged into a single brain value and small intestine and colon were averaged into a single intestine value.

The tissue-specificity index was then recomputed for both species over this nine-tissue panel, using the definition given above: τ was calculated on log2(mean expression + 1), left undefined where the maximum per-tissue mean was below 1, and the enriched tissue was taken as the tissue of maximum mean expression. This nine-tissue τ is specific to the cross-species comparison and is not the eleven-tissue τ reported elsewhere in this descriptor. Medaka and human genes were joined through the one-to-one ortholog table on the human gene symbol, and correspondence was evaluated among orthologs that were tissue-specific (τ ≥ 0.8) in both species.

Because the human values derive from post-mortem tissue processed on a different platform, and because tissue homology between a teleost and a mammal is approximate, the comparison is treated as a descriptive indicator of shared expression patterns rather than as evidence of functional equivalence.

### Interactive data browser

The dataset is accompanied by an interactive browser, the Medaka RNA-seq Dashboard, implemented as a single self-contained HTML file that runs entirely client-side. Both third-party libraries (Plotly.js v2.32.0, MIT; SheetJS CE v0.18.5, Apache-2.0) and the anatomical diagrams are embedded, so the file has no server-side component, no external dependency and no network requirement; library versions are fixed at build time.

The released expression, principal-component and differential-expression tables are loaded locally by the user. Five panels are provided: a tissue browser that maps the expression of a selected gene onto a mid-sagittal diagram of the fish and opens per-tissue detail on click, and that can show a female and a male diagram with the four conditions averaged with equal weight, the two levels of any of the three axes side by side, or a difference map of the log2 ratio between them; a gene-expression panel showing one gene across all eleven tissues for any of the three axes or sex-pooled; a heatmap panel that applies hierarchical clustering to a user-supplied gene list, with a choice of aggregation, scaling, distance metric and linkage; a principal-component panel; and a differential-expression panel giving the tissue × axis summary matrix, which is recounted when the adjusted p-value and log2 fold change thresholds are changed, and the filtered tables, with export to CSV. The bar plots and the principal-component plot can be saved as PNG images, and the heatmap as SVG or PNG. Gene symbols are linked across panels, so a gene found in a differential-expression table can be inspected directly in its expression plot or on the anatomical map. The interface is available in English and Japanese. The anatomical diagrams were drawn by the authors and are released under CC-BY-4.0.

### AI-use statement

Large-language-model assistance (Claude, Anthropic) was used for English editing and for the development of the analysis scripts and the interactive browser; all analyses and the browser were directed and verified by the authors.

### Data Records

All raw sequencing reads have been deposited in the DDBJ Sequence Read Archive (DRA) under BioProject PRJDB45856^36^. Each of the 91 sequenced libraries is registered as a separate BioSample (SAMD02113833–SAMD02113923) and DRA run (DRR1207688– DRR1207778). Per-sample identifiers and attributes (strain, tissue, sex, environmental axis and condition, collection site and date) are given in Supplementary Table S1, and the sample composition is summarized in Table 1. The single library excluded after quality control (RSwMT1) is retained in the deposit and flagged (use = FALSE) so that the complete sequencing output remains available.

Processed data, analysis scripts and the interactive browser are available from Zenodo^37^. The expression matrices, the sample principal-component scores, the tissue-specificity table and the human-ortholog mapping are at the top level of the deposit. The remaining files are organized into de/ (differential-expression tables), de_diagnostics/ (design and filtering summaries of the differential-expression fits), pairwise_comparison/ (the pairwise library comparison shown in Supplementary Fig. S1), tau_cohort/ (tissue-specificity indices by cohort), crossspecies/ (cross-species annotation), qc_strand_mapping/ (strandedness and mapping summaries), metadata/ (sample labels, per-library quality-control values and DDBJ accessions), dashboard/ (the self-contained browser, its user guide and its test), supplementary_tables/, external/ (the NCBI and Human Protein Atlas files read by the scripts), env/ (the software environments) and scripts/. Supplementary Table S2 describes the released files and directories with their formats, contents and column definitions; MANIFEST.tsv in the deposit lists every file with its size and checksum.

Expression is reported for 24,488 genes across the 90 analyzed libraries, as transcripts per million (TPM) and as tximport estimated counts. In the spreadsheet versions of the two matrices, each gene symbol is a hyperlink to the corresponding NCBI gene record; the TPM matrix is also provided as plain text, which is the file read by the dashboard. Differential-expression tables are provided separately for each somatic tissue and each of the three axes (27 tables), and report the shrunken log2 fold change used for the |log2 fold change| > 1 criterion together with the unshrunken maximum-likelihood estimate, so that users can apply either. The tissue-specificity table gives, for each gene, the τ index, the enriched tissue and the mean expression in each of the eleven tissues; human orthologs are supplied as a separate mapping table.

### Technical Validation

#### Read quality

Input RNA was of high integrity (RINe median 9.2, range 7.5–10.0; Supplementary Table S3). After fastp filtering, a median of 99.5% of reads passed quality filtering (range 99.0–99.7%), with a median post-filtering Q30 base rate of 95.4%. Retained libraries comprised a median of 27.3 million read pairs (range 24.8–46.1 million). Per-sample metrics are provided in Supplementary Table S3.

#### Mapping

Reads were quantified against the medaka transcriptome at a median mapping rate of 86.6% (range 76.9–95.1%) across the 90 analyzed libraries, consistent with high-quality, on-target RNA-seq (Table 3). Library strandedness was confirmed directly from the data: salmon inferred the ISR (reverse-stranded, inward) library type for all 90 analyzed libraries, with a median reverse-strand fragment fraction of 0.985 (range 0.966–0.993) and a median compatible-fragment ratio of 1.00, consistent with the dUTP-based Illumina Stranded mRNA Prep chemistry (Supplementary Table S4). Mapping rate varied modestly among tissues (per-tissue medians 80.8–93.2%; highest in liver and ovary, lowest in heart and brain). Per-sample and per-tissue values are provided in Table 3 and Supplementary Table S3.

**Table 3.** Mapping rate by tissue. Median, minimum, and maximum salmon mapping rate (%) per tissue across the 90 analyzed libraries. Mapping is uniformly high (per-tissue medians 80.8– 93.2%), with no tissue showing systematic mapping failure (see Table 1 for sample breakdown).

| **Tissue** | **n libraries** | **Median mapping (%)** | **Min (%)** | **Max (%)** |
|---|---|---|---|---|
| Liver | 9 | 93.16 | 91.3 | 94.76 |
| Ovary | 4 | 92.25 | 90.24 | 93.64 |
| Testis | 4 | 90.17 | 85.67 | 92.06 |
| Intestine | 9 | 88.91 | 85.83 | 91.09 |
| Gill | 9 | 88.02 | 86.05 | 89.46 |
| Muscle | 10 | 87.88 | 80.18 | 95.08 |
| Spleen | 9 | 86.62 | 84.22 | 89.08 |
| Kidney | 9 | 85.48 | 79.07 | 89.34 |
| Eye | 9 | 84.68 | 83.71 | 86.46 |
| Brain | 9 | 82.67 | 80.09 | 86.02 |
| Heart | 9 | 80.81 | 76.91 | 86.55 |
| **All (analyzed)** | **90** | **86.59** | **76.91** | **95.08** |

#### Sample structure and outlier detection

In PCA of the 5,000 most variable genes, samples grouped primarily by tissue rather than by sex or environmental condition; the first two components captured 27.1% and 14.7% of the variance (Fig. 1). Within each tissue, salinity- and season-axis samples intermingled, indicating that tissue identity, not environmental treatment, is the dominant source of transcriptional variation. Pairwise sample correlations showed a clear block-diagonal structure, with libraries of the same tissue forming contiguous high-correlation regions. Certain pairs of tissues off the diagonal also showed high correlation. The initial screen of all 91 collected libraries flagged one mislabeled library: RSwMT1, nominally testis, correlated most strongly with intestine (nearest-neighbor intestine sample RSwMI1, r ≈ 0.94). It was excluded, and Fig. 1 shows the 90 retained libraries.

#### Recovery of tissue-specific expression

A tissue-specificity index (τ) was defined for 21,959 genes (median τ = 0.54; Fig. 2). The number of tissue-enriched genes (τ ≥ 0.8) was highest in brain (1,821), followed by eye (779), gill (730), testis (571) and ovary (458), and lowest in heart (129) and spleen (112), consistent with the known transcriptional complexity of neural and reproductive tissues^35^; the eye and the gill are likewise composed of multiple specialized cell types.

**Fig. 2.**
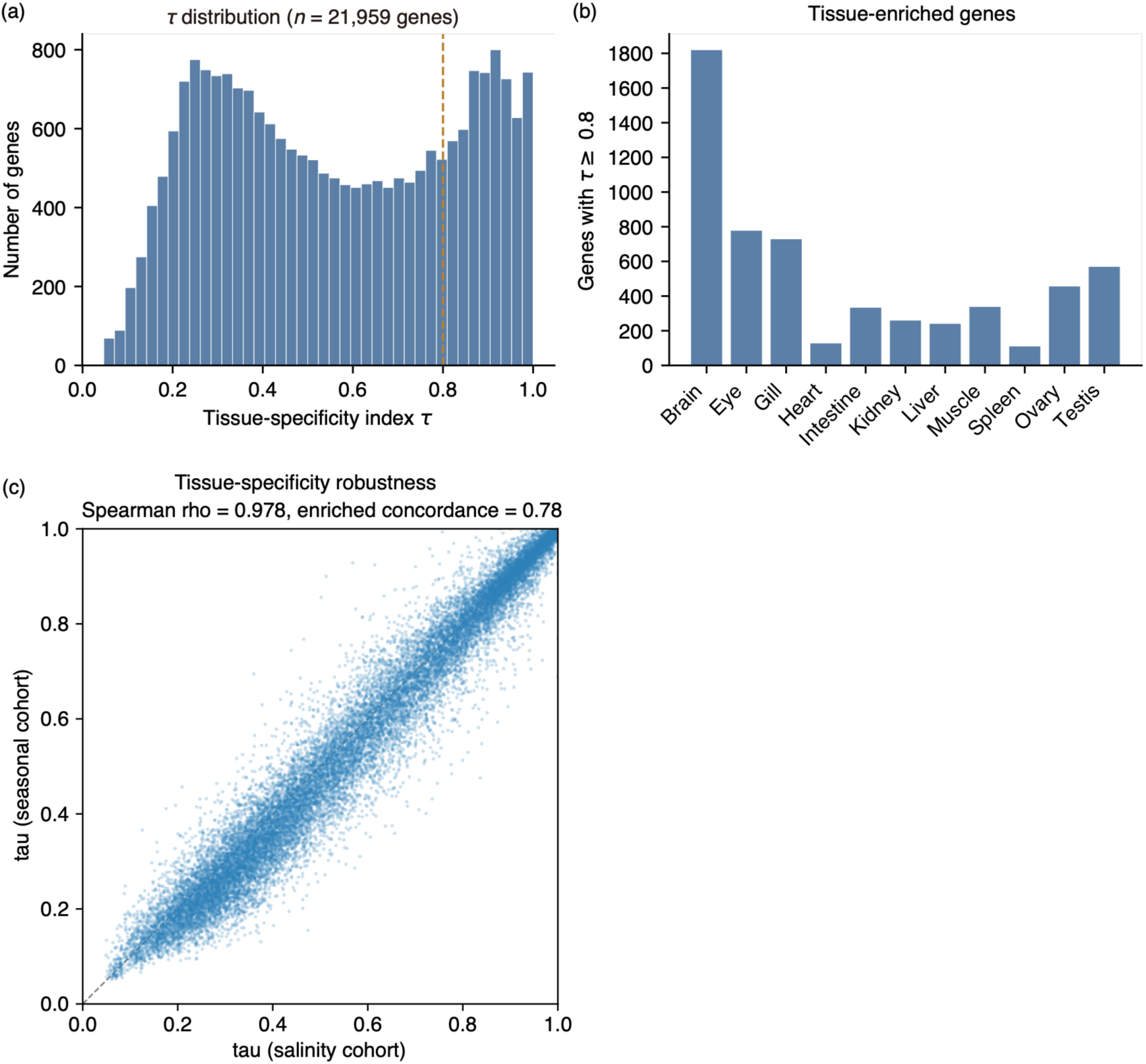
Tissue-specificity. (a) Distribution of the tissue-specificity index τ across genes with defined τ (n = 21,959). (b) Number of tissue-enriched genes (τ ≥ 0.8) per tissue. (c) τ computed independently within the salinity and the seasonal cohorts (49 and 41 libraries) for the 21,616 genes with τ defined in both; the dashed line is the identity (Spearman ρ = 0.978).

To characterize the tissue-restricted fraction of the atlas, we listed the ten genes with the highest τ in each tissue, restricted to genes whose maximum per-tissue mean reached 100 TPM so that the resulting set is usable as a practical marker panel (Supplementary Table S5). Genes were ranked by τ alone; no selection based on prior knowledge was applied. The resulting 110 genes span τ = 0.76–1.00 and 102–23,519 TPM, and include many genes with established tissue-restricted roles in vertebrates: the cardiac sarcomeric genes *myl7*, *vmhc*, *cmlc1* and *csrp3*; the hypothalamic and pituitary hormones *avp*, *oxt* and *prl* together with the vesicular glutamate transporter *slc17a6a* in brain; the proximal-tubule transporters *slc5a2* and *slc34a1b* in kidney; the ionocyte-associated genes *foxi3b*, *atp6v1c2* and *slc4a1b* in gill; the meiotic genes *sycp1*, *smc1b* and *tex12* in testis; *aqp10a* and *mogat2*, which mediate water and lipid absorption in intestine; and *tlx1* in spleen, the transcription factor whose loss causes isolated asplenia in zebrafish^38^. Their recovery without any curation indicates that the atlas reproduces expected tissue identities. Fifty-seven of the 110 genes carry provisional LOC identifiers, of which 38 have descriptive annotations (for example, plasma protease C1 inhibitor-like and coagulation factor VII in liver), so the list also nominates candidate tissue-restricted genes that are not yet formally named; in the ovary, all ten are currently uncharacterized. The spleen had the shallowest tissue-restricted set: only five of its ten top-ranking genes reached τ ≥ 0.8 (median τ = 0.809), consistent with its having the fewest tissue-enriched genes overall (112). Its highest-ranking genes were markers of splenic identity and stroma (*tlx1*, *ccbe1*, *grem2a*, *nppc*) and the thrombopoietic genes *gp1ba* and *mpl*; the latter, though 6–7-fold enriched in spleen, retain secondary expression in kidney, heart and gill, which caps τ.

To confirm that the tissue-specificity index is not driven by pooling the two cohorts, τ was compared between the two cohort-wise estimates (49 and 41 libraries). The two were highly concordant (Spearman ρ = 0.978 across 21,616 genes with τ defined in both), and among tissue-specific genes (τ ≥ 0.8 in both cohorts) 94.3% shared the same enriched tissue. Lower enriched-tissue agreement among all genes (77.9%) is expected for broadly expressed, low-τ genes, whose maximally expressing tissue is labile by definition. The τ index is therefore robust to cohort composition (Fig. 2c).

#### Environmental responsiveness

Per-tissue differential-expression analysis identified candidate condition-associated genes with tissue-dependent patterns on the salinity and seasonal axes (gonads were not tested, having a single replicate per condition). Along the salinity axis, candidate genes meeting the exploratory DE threshold (padj < 0.1 and |log2 fold change| > 1) were most numerous in kidney (1,144 genes) and gill (942), followed by intestine (394) and muscle (393), and were fewest in brain (15). The predominance of kidney and gill is consistent with the organ-specific demands of ion and water balance in these tissues^10–12^. Along the seasonal axis, large numbers of condition-associated candidate genes were detected in every somatic tissue (2,040–3,821 genes; highest in muscle and kidney), consistent with the broad, multi-tissue nature of temperature- and season-associated responses^9,13^. Along the rearing axis, 417–3,576 candidate genes were detected per somatic tissue (highest in muscle, lowest in brain). Summary counts per tissue and axis are provided with the processed data (Fig. 3).

**Fig. 3.**
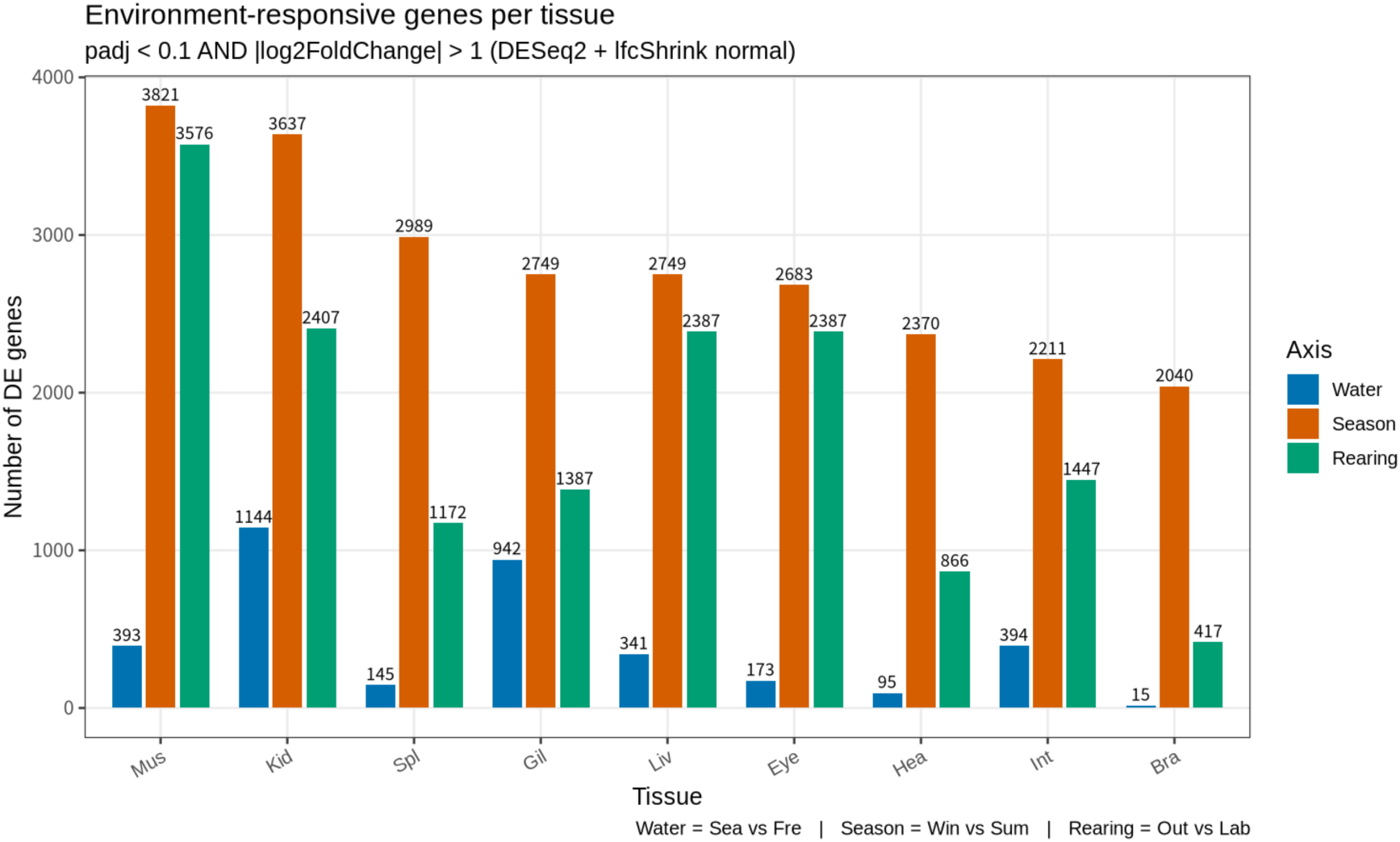
Environmental responsiveness. Number of differentially expressed genes (padj < 0.1 and |log2 fold change| > 1) per tissue for the salinity (seawater versus freshwater), seasonal (winter versus summer) and rearing (summer group versus freshwater group) axes. The salinity and seasonal axes derive from separate cohorts sampled at different facilities, and the rearing axis reuses libraries of both; absolute DE-gene counts should not be compared quantitatively between axes. Counts are for the nine somatic tissues only; gonads (one replicate per condition) were not tested.

Each condition has its own sampling date, so sampling date is confounded with condition on all three axes (Table 2). Sequencing batch differs between the axes. On the salinity axis, the freshwater libraries and the seawater libraries of SwF2 and SwM1 were sequenced in one batch (2024-05-21), and those of SwM5 were sequenced later (2024-11-12). On the seasonal axis, all summer libraries were sequenced in one batch (2024-06-18) and the winter libraries in earlier batches, apart from RWFM1. On the rearing axis, the freshwater and summer libraries were sequenced in different batches. Season and rearing are therefore confounded with sequencing batch, whereas salinity largely is not. We treat the seasonal and rearing differential-expression results as exploratory and do not compare differential-expression gene counts between axes.

#### Batch effect

Two sets of libraries were used to descriptively compare within-condition variations with between-condition differences. For pairs of libraries from the same tissue and sex, we computed the fraction of genes whose DESeq2-normalized counts differed by more than twofold (|log2 ratio| > 1, with a pseudocount of 1), among genes with a mean normalized count of at least 100 in the pair (Supplementary Fig. S1).

The first set is the two winter female muscle libraries, RWFM1 (from WF1) and RWFM2 (from WF2), which share condition and sampling date and differ in individual and in sequencing batch; RWFM1 was sequenced in the same batch as the summer libraries. The fraction for this pair was 0.03, compared with 0.31 for the winter-versus-summer pairs and 0.32 for the rearing pair of the same tissue and sex.

The second set is the two seawater males, SwM1 and SwM5, which share condition, tank and sampling date and differ in individual, in the time between sampling and RNA extraction (several days to several weeks versus about eight months) and in sequencing batch. Across the nine somatic tissues, the fraction for this pair ranged from 0.03 to 0.23 (median 0.14). In males, the winter-versus-summer pair exceeded it in all nine tissues (median ratio 1.9), and the rearing pair exceeded it in eight of nine tissues (median ratio 1.6; the exception was spleen). In addition, the difference between the freshwater male (FwM1) and a seawater male was not consistently larger for SwM5 (later extraction and sequencing batch) than for SwM1 (same batch as FwM1): it was larger in five tissues and smaller in four.

These comparisons are based on single pairs of libraries and are descriptive. In these particular comparisons, within-condition differences were generally smaller than between-condition differences on the seasonal and rearing axes; however, batch effects could not be separated from individual-level variation.

#### Cross-species annotation

Of the 24,488 genes, 9,480 (38.7%) had a one-to-one human ortholog in NCBI gene_orthologs. Conserved orthologs were over-represented among broadly expressed genes: 42.8% of expressed genes, but only 23.7% of tissue-enriched (τ ≥ 0.8) genes, had a one-to-one human ortholog, consistent with the greater evolutionary conservation of broadly expressed genes relative to tissue-restricted ones^39,40^. To validate the human-ortholog annotation released with this dataset, we recomputed τ identically in medaka and the human reference (Human Protein Atlas v25.1) over a shared nine-tissue panel and asked whether tissue-specific orthologs are enriched in the corresponding tissue in both species. Among one-to-one orthologs tissue-specific in both species (τ ≥ 0.8; n = 567), 73.7% showed the same enriched tissue in medaka and human, compared with 25.7% across all orthologs with τ defined in both species. Correspondence was high for testis, skeletal muscle, and liver, which have deeply conserved vertebrate expression programs: it was approximately at the overall level for brain, and lower for ovary, spleen, and kidney. Among the 21 orthologs enriched in medaka ovary, the most frequent human counterpart was testis (12), consistent with the sharing of germ-cell and meiotic genes across the gonads. Estimates for tissues with few tissue-specific orthologs (e.g., spleen, heart) are based on small sample sizes and should be interpreted with caution. This correspondence indicates that the released per-gene ortholog annotation (available at the repository) supports medaka-to-human comparison (Fig. 4).

**Fig. 4.**
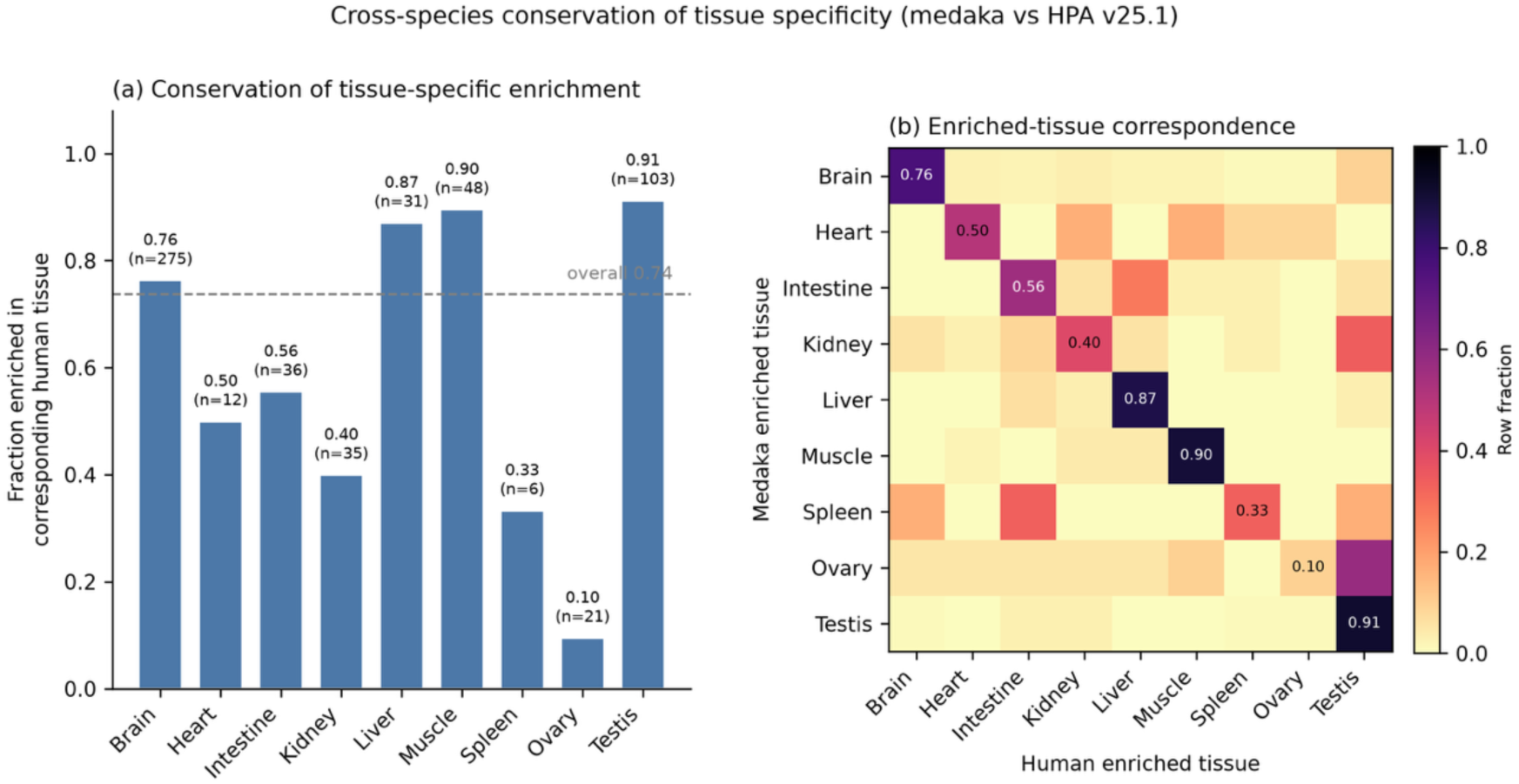
Cross-species conservation of tissue-specific expression (medaka vs Human Protein Atlas v25.1). For 1:1 medaka–human orthologs, the tissue-specificity index (τ) was recomputed identically in both species over a shared nine-tissue panel (eye and gill excluded, lacking a human counterpart). **(a)** Among orthologs tissue-specific in both species (τ ≥ 0.8 in each; n = 567), the fraction enriched in the corresponding human tissue, by medaka enriched tissue (n per tissue indicated); the dashed line is the overall fraction (0.74). **(b)** Row-normalized correspondence between the medaka enriched tissue (rows) and the human enriched tissue (columns) for the same orthologs; diagonal values equal the fractions in (a). Estimates for tissues with few tissue-specific orthologs (e.g. spleen, heart) are based on small samples.

### Usage Notes

The atlas is accompanied by the Medaka RNA-seq Dashboard (Fig. 5), which allows the data to be explored without programming or installation.

**Fig. 5.**
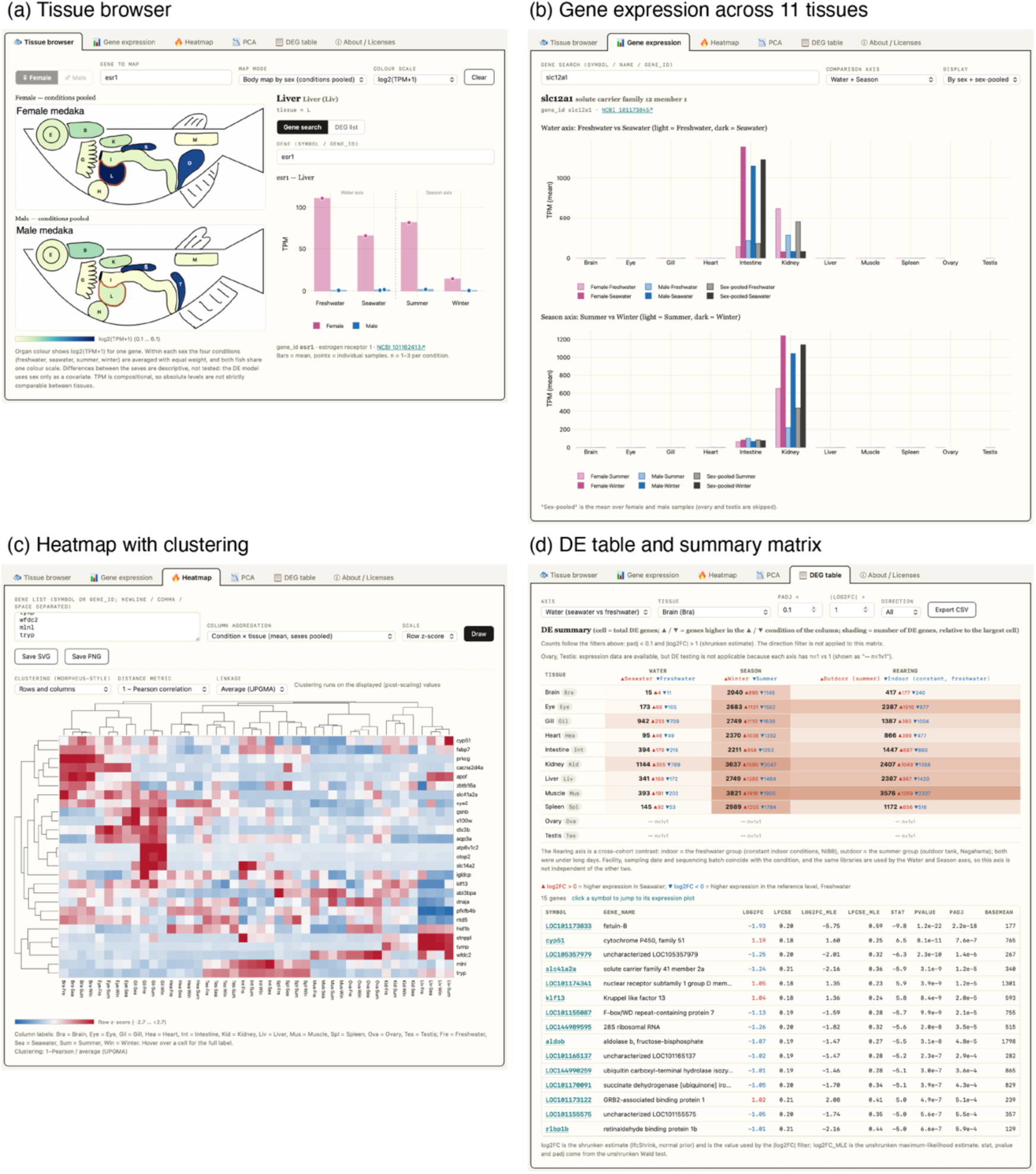
The Medaka RNA-seq Dashboard. The atlas is accompanied by an interactive browser distributed as a single, self-contained HTML file that runs entirely in a web browser without installation, server or network access. **(a)** Tissue browser: a mid-sagittal anatomical diagram doubles as an expression map, coloring every tissue by the expression of a selected gene (here *esr1*, log2(TPM+1), female and male diagrams, with the four conditions averaged with equal weight); clicking a tissue (here liver) opens per-tissue detail, showing expression in the four conditions with bars as means and points as individual samples. **(b)** Gene expression: a selected gene (here *slc12a1*) across all eleven tissues, shown separately for the salinity and seasonal axes and split by sex, with the sex-pooled mean. **(c)** Heatmap: a user-supplied gene list (here the three top-ranked differentially expressed genes of each somatic tissue on the salinity axis) with hierarchical clustering (row z-score; 1 − Pearson distance, average linkage); columns are labeled with the tissue and level codes given in Usage Notes (for example, Kid·Sea, seawater kidney). **(d)** DEG table: the per-tissue × per-axis differential-expression summary matrix for the three axes (gonads marked “— n=1v1”, as they are not tested; counts follow the padj and |log2 fold change| filters) above the filtered gene table for the selected combination (here kidney, salinity axis, padj < 0.1, |log2 fold change| > 1); gene symbols link to the corresponding expression plot, and the filtered set can be exported. In the summary matrix, ▴ and ▾ count the genes with higher expression in the condition marked ▴ or ▾ in the column header.

Five sampling definitions affect interpretation. The brain sample included the pituitary, so pituitary hormone genes (for example *prl* and *gh1*) are represented in the brain profile. The kidney sample included the head kidney, which is the principal hematopoietic organ in teleosts, so the kidney profile contains a substantial immune-cell component. The eye was sampled whole, so lens crystallins are represented alongside retinal genes. Muscle refers to caudal trunk musculature, which in teleosts is dominated by fast (white) fibers. Because the digestive tract comprises multiple functional regions, ‘intestine’ is used as an operational label for the entire tract; this anatomical scope should be considered when interpreting region-specific absorptive markers.

In the released files and in the dashboard, the salinity axis is labeled “Water”, the seasonal axis “Season” and the rearing axis “Rearing.” The two levels of the salinity axis are “Sea” (seawater) and “Fre” (freshwater), those of the seasonal axis are “Win” (winter) and “Sum” (summer), and those of the rearing axis are “Lab” (the freshwater group; shown in the dashboard as “Indoor (constant, freshwater)”) and “Out” (the summer group; shown as “Outdoor (summer)”); samples not informative for a given axis are marked “ns”. Tissues are labeled with three-letter codes: Bra (brain), Eye (eye), Gil (gill), Hea (heart), Int (intestine), Kid (kidney), Liv (liver), Mus (muscle), Spl (spleen), Ova (ovary) and Tes (testis). Differential-expression files follow the pattern de_<axis>_<tissue>.csv.

Interpretive constraints are stated within the interface itself where they apply. In particular, TPM is compositional and is therefore not a quantitative measure for comparing absolute abundance between tissues, so the tissue colors of the tissue browser should be read as a qualitative guide and statistical inference should rest on the DESeq2 results computed from counts; the difference maps are computed within one sex and hence rest on a single replicate per condition, making them descriptive rather than tests; and ovary and testis, having one replicate per condition on each axis, are shown with expression values but are marked as not differential-expression tested. The summary matrix of the differential-expression panel is recounted when the thresholds are changed; counts at adjusted *p*-value thresholds other than 0.1 are descriptive, because the adjusted *p*-values were computed with independent filtering at a 10% false discovery rate.

Libraries were poly(A)-selected, so small-RNA and many non-polyadenylated biotypes (e.g., miRNA, sno/snRNA, tRNA) are not quantitatively captured; values for these features in the TPM, τ, and DE tables should not be interpreted. While count and TPM matrices include these non-polyadenylated features, their quantification may not accurately reflect biological abundance. We recommend restricting analyses to the protein-coding and lncRNA subsets as a default, utilizing the provided annotation columns.

To support translational and comparative work, genes are linked to their human orthologs where available, allowing medaka expression patterns to be related to human genes. Several design features should guide reuse. The salinity and seasonal contrasts were sampled from separate cohorts; comparisons should therefore be made within, not across, axes. The rearing axis is the one defined comparison across the two cohorts. It contrasts the freshwater group with the summer group, which differ in facility, sampling date, sequencing batch and temperature as well as in rearing setting, and it reuses the libraries of the other two axes, so its results are not independent of them. Because ten tissues were taken from each individual, tissues can be compared within individuals using the individual identifier given in Supplementary Table S1. Biological replication per condition is limited (typically one male and one female per condition, with gonads represented by a single sex), so the differential-expression tables are best used to generate hypotheses and prioritize candidate genes rather than as definitive statistical tests, and effect sizes and adjusted p-values should be interpreted accordingly. The tissue-specificity index is relative to the eleven-tissue panel sampled here. Expression is reported against the GCF_053564925.1 annotation; users integrating other datasets should map identifiers accordingly.

## Data Availability

Raw reads: DDBJ DRA, BioProject PRJDB45856. Processed data and the interactive browser: Zenodo DOI: 10.5281/zenodo.23227120. Processed data are released under CC-BY-4.0 and analysis code under the MIT License. The NCBI and Human Protein Atlas files redistributed in the deposit remain under the terms of their providers. The release is versioned (v1.0.0), and the Zenodo record links the dashboard HTML, processed tables, and scripts to this version.

## Code Availability

Analysis scripts (read processing, salmon indexing and quantification, tximport aggregation, PCA and outlier inspection, differential expression, pairwise library comparison, tissue specificity, human ortholog mapping and cross-species comparison) and the dashboard with its browser test are available at Zenodo DOI: 10.5281/zenodo.23227120. Software versions: fastp v0.23.2, salmon v1.10.3, R v4.4.2, tximport v1.34.0, DESeq2 v1.46.0, Python v3.11.15, numpy v2.4.6, pandas v3.0.3, matplotlib v3.11.0 and scipy v1.17.1. The scripts are arranged to run on a single Linux machine, in two conda environments defined in the deposit (one for read processing and the R analyses, one for the Python analyses). The README of the deposit gives the order of execution, the inputs and outputs of each step, and the steps that were re-run in this arrangement to confirm that they reproduce the released files.

## Author Contributions

TK, YT and KN conceived and designed the study. YT and KN provided fish. YT, AT and KN performed experiments. TK performed data analysis. TK wrote the first draft of the manuscript with YT and KN making significant contributions to editing. All authors agree to be held accountable for the content therein and approve the final version of the manuscript.

## Acknowledgements

We thank NBRP Medaka (https://shigen.nig.ac.jp/medaka/) for providing the d-rR/TOKYO strain (Strain ID: MT837), and Dr. Eitaro Sawayama (Nihon University) and Dr. Tomoya Nakayama (Nagoya University) for discussions on the treatment conditions of medaka. We also thank Ms. Erina Fukunaga for technical assistance with nucleic acid extraction.

