## Supplementary Figure 1 for "A multi-tissue RNA-seq atlas of adult medaka (*Oryzias latipes*) under salinity and winter–summer sampling contrasts"

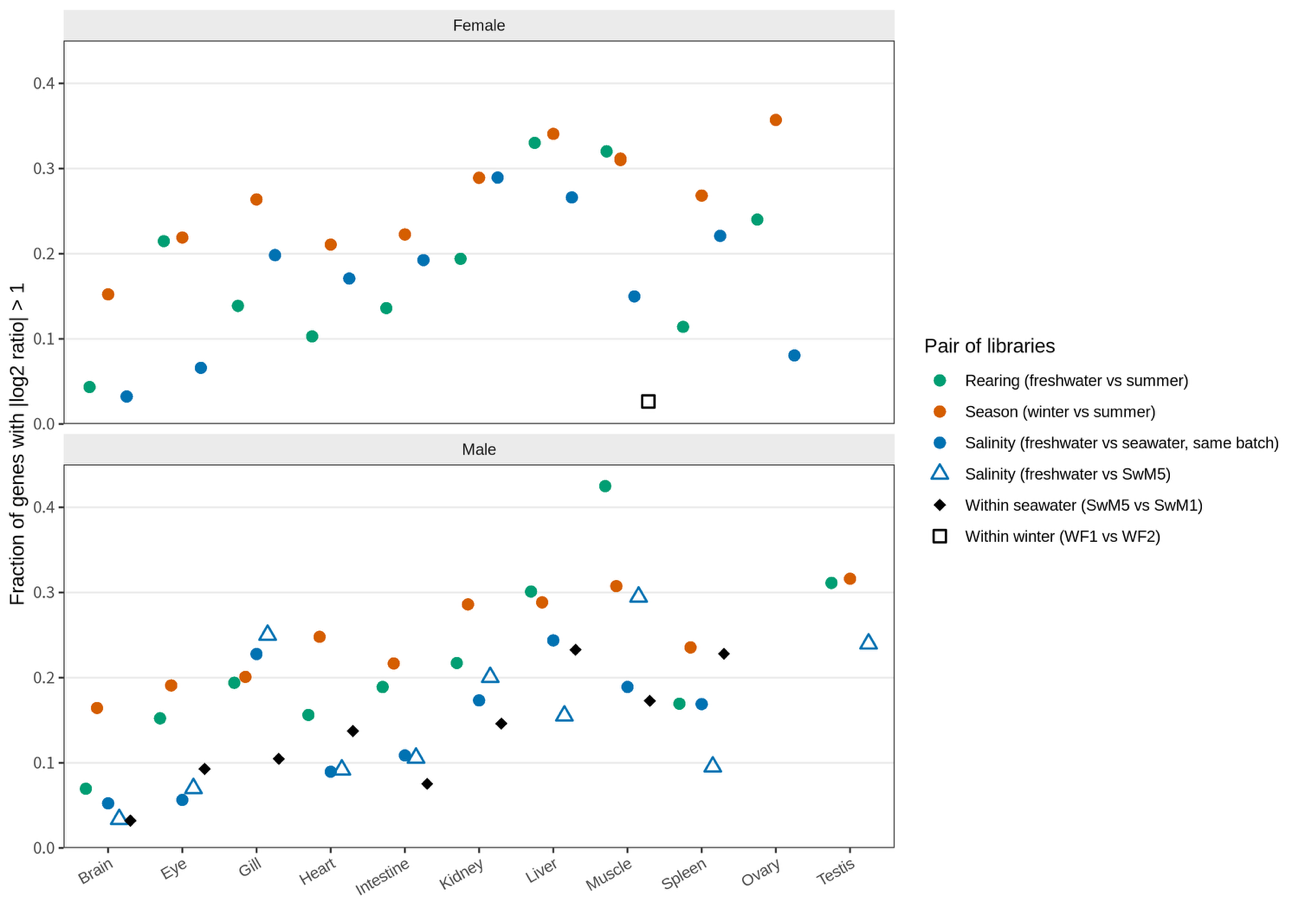

**Supplementary Fig. S1.** Differences between single pairs of libraries within tissue. For each pair of libraries from the same tissue and sex, the fraction of genes with |log2 ratio| > 1 on DESeq2-normalized counts (pseudocount 1; genes with a pair-mean normalized count of at least 100). Pair classes: rearing (freshwater versus summer), season (winter versus summer), salinity (freshwater versus SwF2 or SwM1; same sequencing batch), salinity with SwM5 (different batch), within seawater (SwM5 versus SwM1) and within winter (RWFM1 versus RWFM2). Each point is one pair; females and males are shown in the upper and lower panels. Ovary and testis are included, although they were not tested for differential expression. For testis, the only salinity pair is that with SwM5, because RSwMT1 was excluded. For female muscle, two season pairs are plotted (RWFM1 and RWFM2, each versus the summer female) and overlap.
